# Histology-guided 3D virtual staining of microCT-imaged lung tissue via deep learning

**DOI:** 10.1101/2025.10.02.678959

**Authors:** Cristina Almagro-Pérez, Niccolò Peruzzi, Csaba Galambos, Andrew H. Song, Hans Brunnström, Kinga I. Gawlik, Marco Stampanoni, Karin Tran-Lundmark, Goran Lovric

**Affiliations:** Institute for Biomedical Engineering, ETH Zurich, 8092, Zurich, Switzerland; Swiss Light Source, Paul Scherrer Institut, 5232 Villigen, Switzerland; Department of Experimental Medical Science Lund University, 22 184 Lund, Sweden; Department of Pathology and Laboratory Medicine, University of Colorado Anschutz School of Medicine, Aurora, USA; Department of Pathology, Mass General Brigham, Boston, MA, USA; Division of Pathology, Department of Clinical Sciences Lund, Lund University, SE-22185 Lund, Sweden; Wallenberg Center for Molecular Medicine, Lund University, 22 184 Lund, Sweden

## Abstract

Histologically stained tissue sections are considered the gold standard for studying microscopic anatomy and diagnosing disease in clinical practice. However, the processes of sectioning and staining are laborious, and the overall method relies on two-dimensional (2D) analysis. In contrast, X-ray-based virtual histology offers the advantage of virtual sectioning while retaining the full three-dimensional (3D) volumetric representation of the tissue. Nevertheless, its grayscale nature has prevented it to be readily utilized by pathologists who are accustomed to conventional histological stains. In this work, we present a histology-guided enhancement platform that can integrate the 3D information provided by synchrotron radiation phase-contrast microCT with the rich visual features characteristic of histological stains. We introduce a multi-stage microCT-histology co-registration method combined with a virtual staining deep neural network and demonstrate successful virtual histological staining of microCT human and mouse lung tissue that closely resembles standard histology. We evaluate our strategy on multiple histological stains and apply it to identify 3D collagen-based remodeling of pulmonary arteries in patients with pulmonary hypertension. Overall, this innovative enhancement pipeline has the potential to aid in the incorporation of microCT into clinical practice, and advance non-destructive 3D pathology for improved diagnostic efficiency and accuracy.

## INTRODUCTION

Histopathology has long been the gold standard tool in research and pathologic diagnostic practice to study biological tissue at a microscopic level. Histology-based practice comprises diverse sample preparation methods and various histochemical and immunohistochemical stains, allowing the visual identification of multiple cell types and tissue structures ^1,2^. However, the overall process of sectioning, staining, and imaging is time-consuming, destructive to the tissue, and it provides information only in two dimensions (2D). For context, a sample obtained upon resection has a typical depth of a few millimeters while the 2D histological sections are 3-5 µm in thickness, representing less than 0.1% of the entire specimen.

Three-dimensional (3D) pathology involves a change of paradigm where the entire tissue specimen is analyzed, and clinical decisions are based upon the 3D morphology. Recent studies have highlighted the critical need for 3D histopathological analysis, offering new insights into cancer risk assessment ^3^, tumor heterogeneity ^4–6^, and tumoral vascular and neural structures. However, common existing techniques for conducting 3D histopathological analysis, involve labor-intensive sample preparation processes or are limited to submillimeter-thick samples ^7^. In the recent past, with the steady increase of X-ray source and detector technology, X-ray-based virtual histology by means of micro-computed tomography has started to gain increasing popularity in studying large biological tissue specimens in 3D and at spatial resolutions down to the micrometer scale ^8–12^. In particular, the utilization of phase-contrast µCT (PCµCT) at both synchrotron ^13^ and lab-based X-ray sources ^14,15^ has allowed for unprecedented soft-tissue discrimination and is currently delivering promising results toward improved diagnostics ^16,17^. Despite this progress, a significant challenge that currently limits clinical adoption is its single-channel (grayscale) nature, which restricts its ability to produce images with sufficient contrast for specificity to match conventional histopathological stains.

Over the past few years, the advent of deep learning techniques has shown potential for virtual staining of 2D histology, digitizing the staining process and offering fast, inexpensive, and accurate alternatives to standard histochemical staining ^18,19^. These have shown promise in both virtual staining of unstained slides ^20,21^ and stain transfer between conventional Hematoxylin and Eosin (H&E) and immunohistochemistry ^22,23^. In the realm of 3D pathology, H&E virtual staining has been successfully integrated with holotomography imaging technology to visualize thin tissue layers (∼ 50 µm) ^7^ as well as non-invasive optical coherence tomography ^24^. While recent studies have explored the combination of histology and PCµCT based on correlative image analysis ^25^, the question on how the specificity of PCµCT images could be improved by using virtual histological staining, remains unresolved.

In this work, we introduce a virtual histological staining framework for thick (*>*2 mm) tissue samples imaged with PCµCT which we call VISTACT, **VI**rtual **STA**ining of micro-**C**omputed **T**omography. VISTACT integrates a 2D-3D registration strategy for micron-wise alignment between PCµCT and histology, with a supervised deep learning approach that learns the conversion between PCµCT and (registered) histological stains, enabling virtual histological staining of PCµCT imaged tissue samples. We apply VISTACT to human lung tissue of patients with pulmonary hypertension (PH) to obtain Elastin van Gieson (EvG)-stained 3D tissue images that further enable the localization and quantification of PH vascular remodeled regions. A distinguishing feature of VISTACT is its ability to adapt to different histological stains and tissue types. To highlight this versatility, we also demonstrate it for H&E virtual staining of submicron (0.88 µm/pixel) PCµCT image data of mouse lung tissue.

## RESULTS

The overall workflow of VISTACT is depicted in Fig. 1. To develop a platform for virtual histological staining of PCµCT volumes, we initially collected four human lung samples (designated S1-S4) from patients transplanted because of PH. For each lung resection, we acquired high-resolution PCµCT images of certain areas of interest (∼ 2-3 per patient), and obtained several histological sections at different depth levels which were stained with EvG staining to highlight PH vascular remodeling (Fig. 1A). In the following, we present the two main steps required for our approach. First, the registration of 2D whole-slide histology images to the 3D volumetric PCµCT scans to obtain pixel-wise alignment (Fig. 1B). This is followed by the training of a deep neural virtual staining network using the PCµCT-histology image pairs as input-output training examples (Fig. 1C). We then discuss how our approach facilitates 3D sample analysis and demonstrate its capability to be extended to additional histological stains.

**Figure 1:**
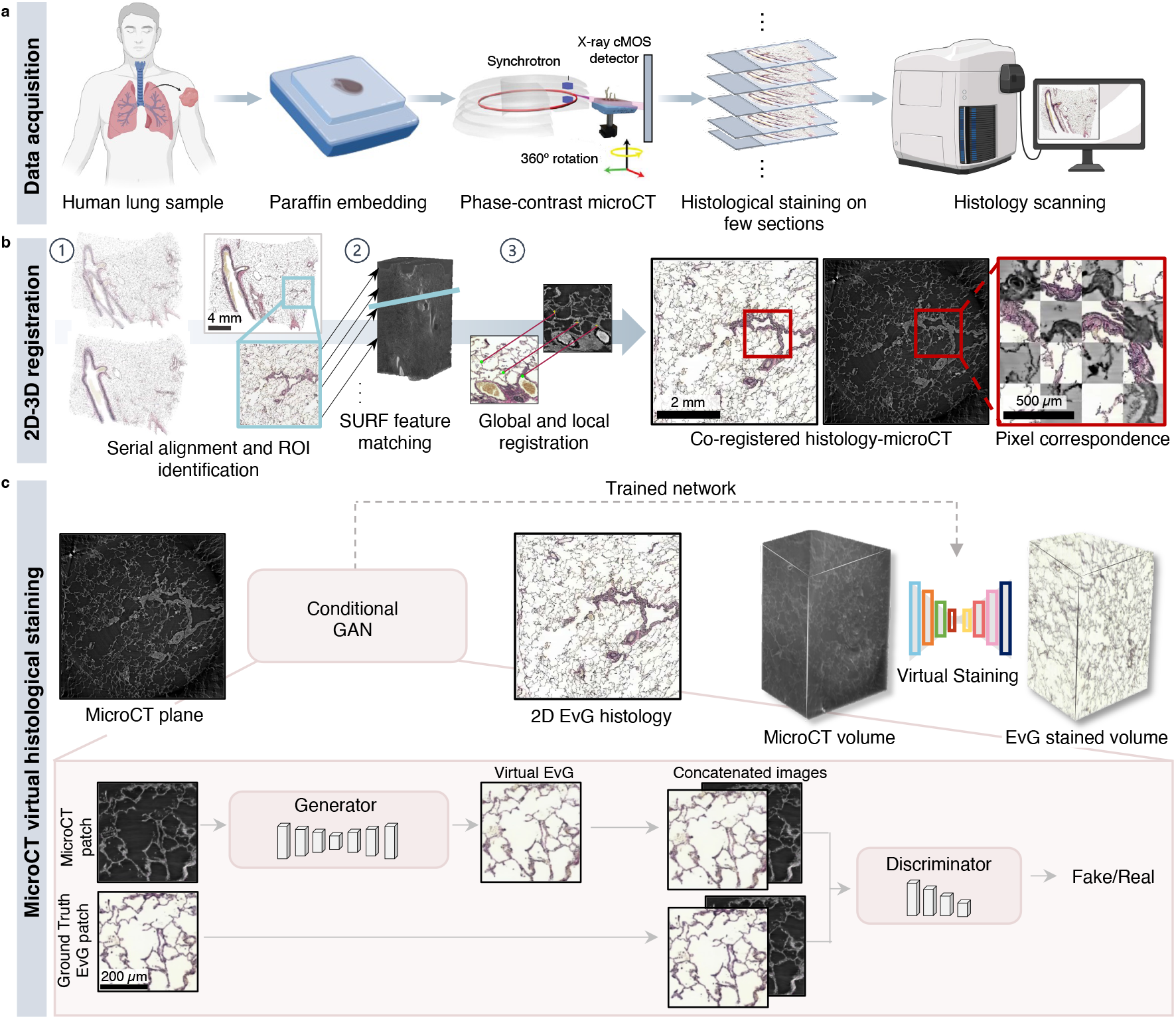
VISTACT: Computational workflow for deep learning-based virtual histological staining of microCT images. **a**. Areas of interest within formalin-fixed paraffin-embedded human lung tissues were subjected to synchrotronbased phase contrast microcomputed tomography. The tissue samples were then serially sectioned, and equally spaced histological sections underwent Elastin van Gieson (EvG) staining and scanning. **b**. The 2D histological sections were matched and aligned to the 3D volumetric microCT images using a three-stage registration pipeline: (1) region-of-interest (ROI) localization within the aligned whole slide images, (2) feature-based matching to find the corresponding microCT oblique plane, (3) final feature-based nonlinear registration of the microCT-histology image pairs to achieve pixel-wise alignment. **c**. The registered microCT-histology images were used as input-output pairs to train a conditional Generative Adversarial Network (cGAN). The trained network was then utilized to virtually stain all microCT axial slices and generate a 3D histological-like volume.

### 2D-3D co-registration for PCµCT-histology alignment

To enable virtual histology based on PCµCT, each PCµCT image needs to be registered to its corresponding histology with the highest accuracy as possible such that virtual staining can occur with a conditional Generative Adversarial Network that maps inputs (PCµCT) to registered outputs (histology). In addition to being precise, the co-registration strategy should require minimal manual effort, such that a large training set of paired PCµCT and histology images can easily be generated. While co-registration is relatively simple when the two imaging modalities are two-dimensional, 2D-3D registration is a much more challenging problem, that requires not only local alignment but this should be preceded by the accurate identification of the corresponding 2D plane within the volumetric image. Previous strategies for PCµCT-histology registration have typically relied on manual identification of the 2D plane ^25^, which is not scalable for registering large numbers of samples required for neural network training, or have been optimized for scenarios where the field-of-view (FOV) is the same across modalities ^26^, a condition that does not apply in our case.

For devising a 2D-3D registration pipeline for PCµCT-histology registration compatible with FOVs of different sizes, we found that accurate image alignment is only possible by a multistage pipeline (Fig. 1B and Fig. S1). The proposed approach consists of (1) serial histological section alignment and approximate identification of the axial location of the PCµCT volumes within the top whole slide image (WSI) for every patient, (2) landmark feature matching to identify the corresponding microCT plane to each histologic section ^27^, and (3) global and local feature-based registration for precise pixel-wise alignment (refer to *Methods* section for details). This approach, for example, successfully aligned 8 EvG-stained histological sections of sample S1 to one of the PCµCT volumes of the same sample in approximately 1 hour (Fig. S2C, breakdown of the time required for each registration step).

We successfully registered 40 histological regions from patients S1-S4 to their corresponding PCµCT volumes. We observed that the Euclidean PCµCT plane corresponding to a histological patch was on average 2^*°*^ tilted with respect to the *Z*-axis. We then evaluated the quality of the alignment once the corresponding PCµCT plane to a given histology image had been found. The Target Registration Error (TRE) was calculated and compared to Elastix ^28^, a widely used medical image registration toolbox. Our method achieved a lower error, with an average of 0.02 ± 0.03 mm distance compared to the 0.23 ± 0.10 mm obtained with Elastix, primarily due to the fact that our registration process explicitly accommodates minor differences in field-of-view, an aspect not addressed by Elastix.

In addition, the results remained consistent when the pipeline was assessed using an additional H&E-stained human lung sample and a H&E-stained mouse heart and lung sample (Fig. S2), demonstrating the robustness of the proposed registration pipeline across samples from different tissue types and histological stains. Through visual inspection, we confirmed micron-level alignment across lung structures including alveoli, bronchioles, respiratory airways and accompanying vessels (Fig. S2A).

Once the histology-PCµCT image pairs were aligned, only image patches (512 x 512 pixels) with high registration accuracy were considered for training (see *Methods* section for details). This step proved to be crucial for PCµCT-histology translation, as the network must capture the genuine staining signal rather than the perturbations caused by misalignments between both imaging modalities. The curated dataset of registered patches was ultimately split into training (80%) and test (20%) sets for training and validation of the virtual staining network.

### Virtual staining of volumetric lung PCµCT image data

The second stage of the proposed pipeline involves a deep neural network to virtually stain PCµCT images as EvG-stained histological sections (Fig. 1C). Here, we propose to adopt *pix2pix* ^29^, a general-purpose image-to-image translation model, which is based on a conditional Generative Adversarial Network (cGAN). This network consists of a generator and a discriminator, which are adversarially trained to map PCµCT images to histology images (Fig. S3). While the generator creates the virtual histologically stained images from input PCµCT images, the discriminator’s role is to distinguish the created image from the ground truth images when provided with chemically stained images.

Using the registered pairs of PCµCT and histology images, the model was given as input the PCµCT images and trained to predict the virtually stained version. Once trained, it was capable of staining, for example, a PCµCT volume of 4.2 mm x 4.2 mm x 1.4 mm, consisting of 840 axial slices, to a resolution of 1.63 µm/pixel in 39 minutes. Visual inspection of the selected cGAN approach revealed that it correctly assigns the desired coloration to the distinct lung structures (Fig. 2), resulting in sharp and EvG-like realistic images. As observed in Fig. 2, the network correctly stains red blood cells contained within the pulmonary capillaries in yellow (Fig. 2e), and provides a pink colorization of the collagen in the external layer of pulmonary airways and a black hue for the lining of the ciliated pseudostratified columnar epithelium in the airways’ internal layer (Fig. 2h). Additionally, the network successfully learned the characteristic EvG colorization of alveolar tissue and collagen, despite these structures exhibit similar PCµCT intensities (Fig. 2b, 2h). It is also noteworthy that the network accurately refrained from staining so-called ring artifacts ^30^, which are commonly found in PCµCT images (some of which are visible in the background of images **a**, and **d**, in Fig. 2). This capability can be attributed to their absence in the corresponding histology counterparts, suggesting the network’s robust adaptability to the unique features of the histological context.

**Figure 2:**
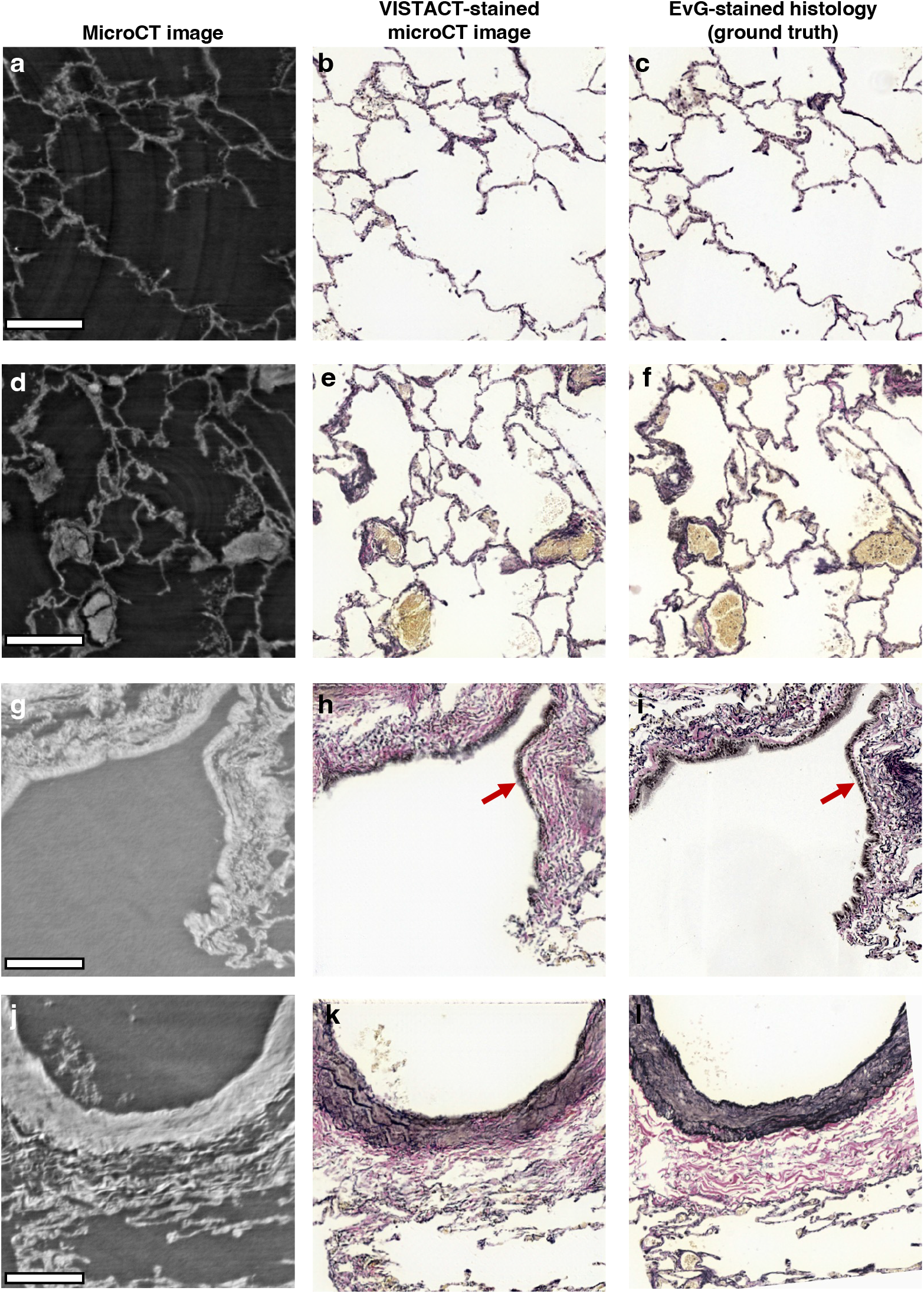
Virtual staining with VISTACT generates EvG-stained-like histology images. **a, d, g, j**. Phase-contrast microCT images of human lung tissue used as input into the virtual histological network. **b, e, h,k** Virtually EvG-stained microCT images. **c, f, i, l**. Bright-field images of EvG-stained histological sections (ground truth). The comparison of **b** and **c** and **e** and **f** demonstrates proper staining of alveoli and correct yellow hue colorization of blood within lung capillaries, respectively. Similarly, in **h** and **i**, the virtual and physical EvG-stained images show correct black colorization of the respiratory epithelial cell nuclei within respiratory airways (red arrows). Correct colorization of blood vessel wall (dark purple) and surrounding collagen tissue (pink) is shown in **j** and **k**. All scalebars are 200 µm.

Prior to the selection of this architecture, other deep learning approaches were investigated. In particular, a Generative Adversarial Network (GAN) previously employed in virtual staining of autofluorescence images was implemented ^20^. However, network convergence was not achieved using this architecture unless the images were subjected to strong preprocessing intended to bring both imaging modalities closer together (Table S1, Fig. S4). In a similar manner and inspired by previous works on image-to-image translation within the medical domain ^31,32^, we also tested a fully convolutional network (FCN) to pose the colorization of PCµCT images as a purely regression problem. However, visual inspection showed that while correct colorization could in this case be achieved in certain lung structures, the images were vastly smoothed, defeating the purpose of PCµCT virtual enhancement (Fig. S5).

In addition to the visual agreement, we evaluated the models using common metrics in virtual staining applications, including the Structural Similarity Index Measure (SSIM), Peak Signal-to-Noise Ratio (PSNR) and Mean Square Error (MSE). While these metrics are widely used, it is important to recognize their limitations, as relying solely on them may yield counterintuitive results; they do not always reflect image quality as perceived by human observers. For this reason, we also incorporate the Learned Perceptual Image Patch Similarity (LPIPs) measure ^33^, which has been reported to outperform previous metrics in terms of closeness to human perception of image quality. The average of these metrics computed on the test set was 0.67 ± 0.12, 17.50 ± 1.63, 0.02 ± 0.007 and 0.25 ± 0.07, respectively. Fig. 4 shows the distribution of these metrics across the patches in the test set. Furthermore, the quantitative comparison of FCN to cGAN based on LPIPs, revealed the superior performance of the cGAN model (0.25 vs. 0.30) for PCµCT virtual histological staining (Table S2).

Notably, by applying the virtual histological staining slice by slice, we can generate entire virtual histological volumes from corresponding PCµCT volumetric images (Figure 3, Movies S1, S2). The capability to create such volumetric representations not only reduce the bias associated to histological sectioning, where only a few sections from an entire specimen are analyzed, but also enables to holistically analyze and examine pathologic changes in their inherent 3D environment.

**Figure 3:**
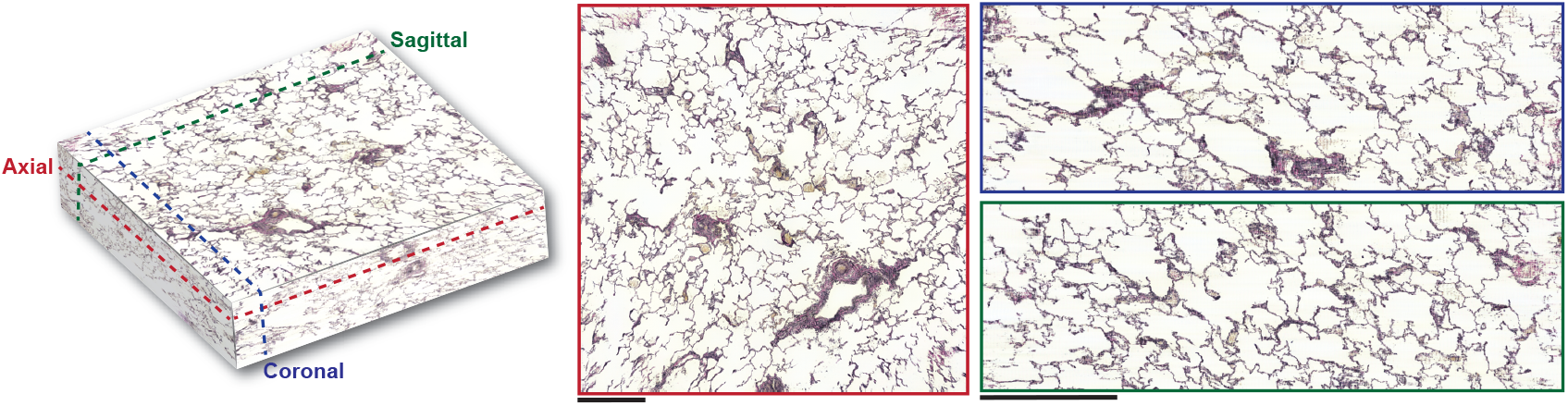
VISTACT generates large millimeter-thick virtually stained volumes. A 3D virtual EvG volume generated slice by slice by VISTACT from a PCµCT-imaged human lung sample. The volume has dimensions 4.2 × 4.2 × 1.4 *mm*^3^. Different regions of the volume can be viewed by slicing the volume across distinct cross-sectional planes for diagnostic purposes.

**Figure 4:**
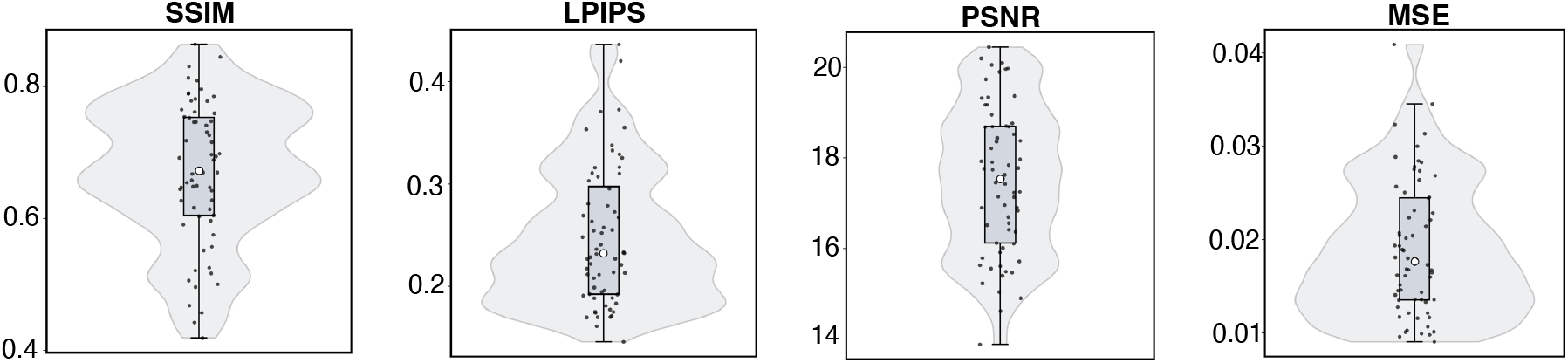
Quantitative evaluation of histological virtual staining with VISTACT. Distribution of SSIM, LPIPs, PSNR, and MSE values computed from 60 patches in the test set. In the box plots, the white circle at the center of the box represents the median, the ends of the box represent the first and third quartiles, and the black dots represent the value for each test patch.

### Unveiling PH vascular remodeled regions in virtually stained PCµCT volumes

Besides facilitating the visualization of the different lung structures, virtual histological staining of PCµCT volumes also enables the subsequent segmentation of critical components of PH. Collagen, which accumulates within remodeled arteries of PH, was segmented from the virtually EvG-stained microCT volumes using color deconvolution (see *Methods* for details). Furthermore, projecting the 3D collagen segmentation masks onto the axial, sagittal, and coronal planes enabled the automatic identification of PH-associated vascular remodeling. As illustrated in Fig. 5, areas of high collagen accumulation corresponded to occluded and dilated vessels resulting from the pathologic processes leading to PH. This illustrates the potential role of 3D virtual staining of PCµCT volumes followed by image processing techniques in identifying disease biomarkers to assist pathologists in enhancing the diagnostic process. Finally, for benchmarking purposes, our overall approach was compared to a standard supervised semi-automatic segmentation on the original (unstained) PCµCT images, implemented in Ilastik ^34^. As shown in Table S3, our proposed approach consistently outperforms the standard method when both approaches are compared to collagen segmentation masks generated from physically stained EvG histological sections; this is likely because Ilastik is limited to features based on microCT intensity and its performance depends on the specific user annotations.

**Figure 5:**
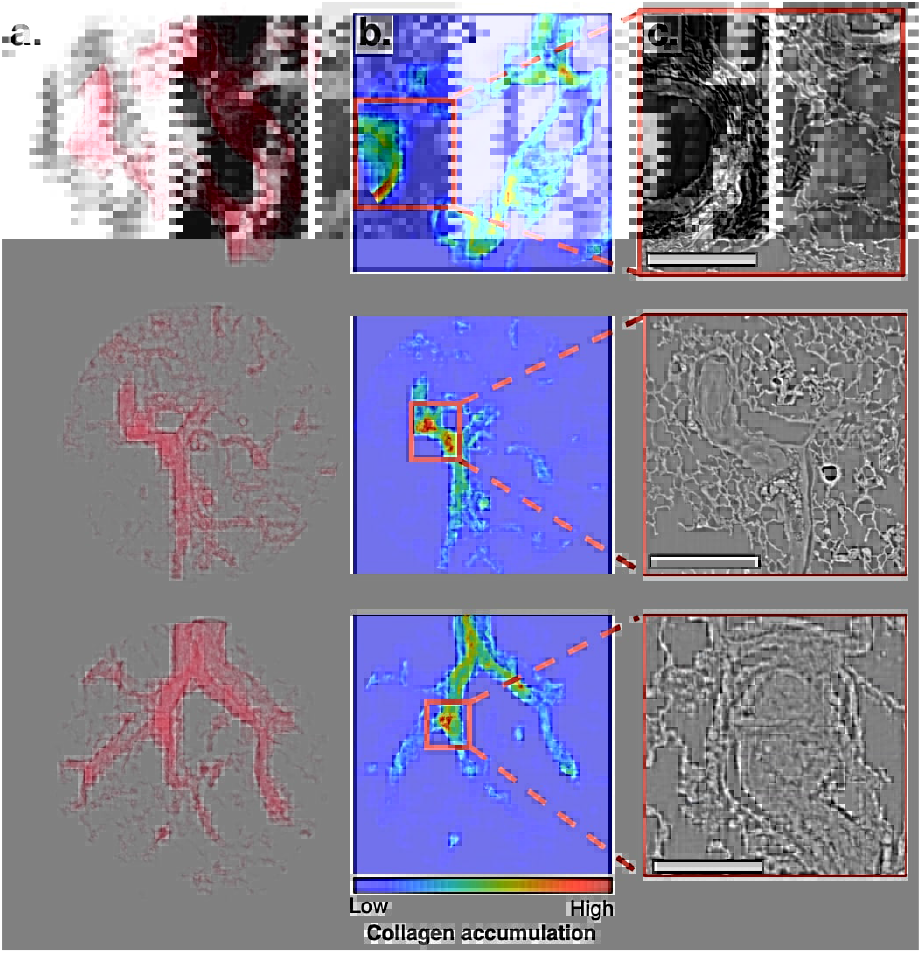
Collagen mask projection facilitates the detection of pulmonary hypertension-related vascular remodeling. Each row represents a separate human lung sample. **a**. 3D collagen segmentation masks obtained from the virtually EvG-stained microCT volumes. **b**. Axial collagen projections. Regions with notable collagen accumulation (shown in red) correspond to vessels with a thickened collagen layer (top), as well as completely obstructed vessels (middle and bottom). **c**. Close-up views of microCT images corresponding to the areas with vascular remodeling. All scalebars are 500 µm.

### H&E-virtual staining of volumetric PCµCT image data

While EvG stain provides unique specific information for the analysis of PH and is widely used for this application ^35,36^, H&E stain is the gold standard in diagnostic pathology practice. As a proof-of-concept study, we tested the proposed pipeline for the task of H&E-histology-guided PCµCT virtual staining. In contrast to the EvG stain, which primarily imparts colorization at the tissue level (i.e. highlighting collagen in pink and elastin in dark purple/black), H&E staining emphasizes structures at the cellular level. The basic hematoxylin component in H&E results in the staining of cell nuclei in blue/purple, contrasting with the pink colorization of the cell cytoplasm and extracellular matrix provided by the acidic eosin. For this reason, higher resolution PCµCT data was acquired for the training of this model. Specifically, a mouse specimen containing heart and lung tissue was imaged with PCµCT at 0.88 µm/voxel resolution and subsequently serially sectioned and stained with H&E (see *Methods* for details).

PCµCT and H&E-stained histologies were aligned with our VISTACT co-registration approach and a separate cGAN virtual staining model was trained for PCµCT-to-H&E conversion (Table S4). As illustrated in Fig. 6, we found strong visual agreement between the H&E-virtually stained PCµCT images and the bright-field images of the same samples that were captured after the histological staining process. While large and bright nuclei are already captured and stained at the current resolution, a higher PCµCT resolution of the scanned volumetric images, will be more suitable to reveal even finer histological details.

**Figure 6:**
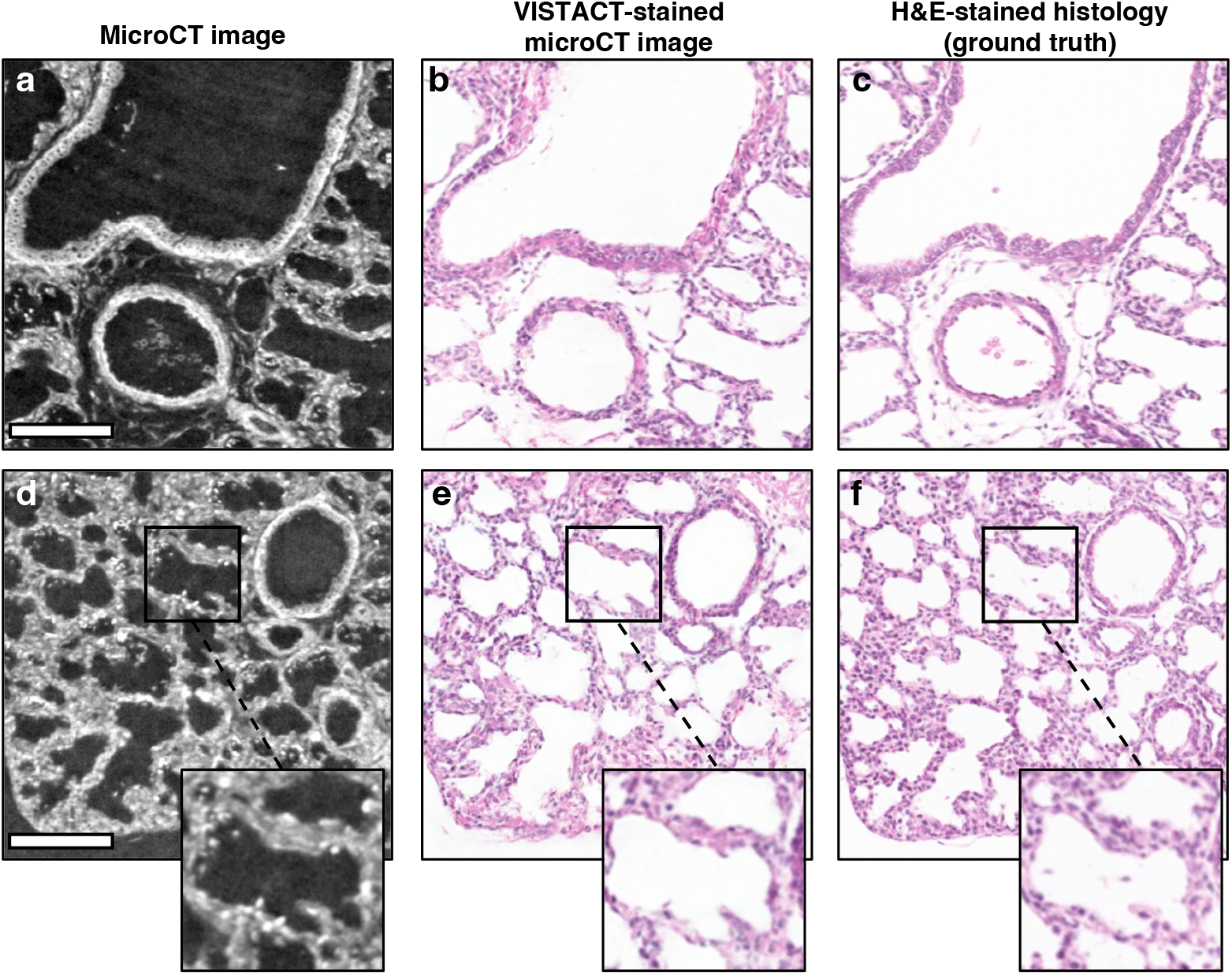
H&E virtual staining of a microCT-imaged mouse lung/heart sample. **a, d, g**: Phase-contrast microCT images at 0.88 µm/pixel are used as input of the virtual histological staining network. **b, e, h**: Virtually H&E-stained microCT images (VISTACT output). **c, f, i**: H&E-stained tissue of the same region captured using a slide scanner with bright field settings. Ground truth for the virtual staining framework. Red arrows highlight examples of stained nuclei. All scalebars are 100 µm.

## DISCUSSION

We introduce a volumetric virtual histological staining framework, VISTACT, which enables the mapping of phase-contrast µCT (PCµCT) images to histological stains appearances, producing high-resolution millimeter-thick 3D samples of human tissue with the conventional color schemes that diagnostic pathologists are trained to analyze. As an initial step in virtually staining PCµCT volumes, accurate registration of PCµCT and histology images was essential. Our proposed multi-stage registration approach eliminates the need for manual PCµCT plane identification and adapts to images with varying fields of view with minimal human input. This enables the rapid and scalable generation of large datasets of paired PCµCT and histology images, which are necessary for training deep neural networks for virtual staining.

The second step toward histology-guided microCT virtual staining involved the selection and training of a virtual staining network. We demonstrated that carefully training a cGAN architecture with precisely registered image pairs, can generate accurate PCµCT images that thoroughly resemble reference histology. In addition to providing a histology-like appearance, that makes 3D pathology volumes available for pathologists and researchers to visually inspect and interpret, the virtually stained PCµCT volumes can facilitate the subsequent segmentation of biological components using conventional segmentation strategies ^37,38^. This has the potential to assist in a clinical setting by highlighting critical diseased regions in 3D. As a proof of concept, we showed that the 3D distribution of collagen obtained from virtually EvG-stained microCT volumes, enables the automatic identification of pulmonary vascular thickening and obstruction, which serve as clear indicators of advanced PH ^39^. We envision that the proposed framework holds significant potential for further addressing a range of pathological challenges beyond PH, including identification of tumor-infiltrating lymphocytes, tumor margins, as well as facilitate 3D studies of intratumoral heterogeneity.

The versatility of VISTACT is also noteworthy. In this work, we demonstrate the flexibility of our pipeline by successfully applying it to multiple histological stains (H&E and EvG), utilizing data acquired from different synchrotron imaging facilities (specifically, the TOMCAT beamline in Switzerland and the ANATOMIX beamline in France), and analyzing specimens from various species, including human lung as well as mouse lung and heart. This adaptability stems from a universally compatible process for sample preparation, imaging, and network training, underscoring the reliability and repeatability of the VISTACT pipeline across varied experimental settings.

In this study, the PCµCT volumes consisted of up to 1,600 axial slices, yet only about 8 histological sections per sample were stained for model training. This highlights the potential to significantly reduce both staining effort and time, while also enabling non-destructive 3D analysis of biological tissues. In future work, expanding the availability of larger patient cohorts could ultimately enable on-the-fly virtual staining of PCµCT volumeseliminating the need for any serial histological sections in new patient specimens.

A limitation of virtual histological staining is the information present in PCµCT images compared to histology, due to the resolution and contrast mechanisms. The resolution of PCµCT images determines the smallest features that VISTACT can accurately reconstruct. For this reason, in this work we started by focusing on EvG staining, which primarily imparts colorization protein abundance at the tissue level, staining collagen layers in pink, elastin in purple/black, respiratory epithelial cell nuclei in black and blood in yellow. We also showed that with sufficient high resolution (0.88 µm/pixel), H&E staining of large and bright nuclei is possible. With the continued development of PCµCT technology with higher imaging resolutions, it is reasonable to expect that even subcellular histological details will be captured and resolved.

In conclusion, we believe that the combination of VISTACT and PC*µ*CT will become a revolutionary tool in diagnostic pathologic practice. We further envision the use of this platform to aid in novel discoveries and understanding of the role microanatomical structures play in disease by examining tissue samples in 3D. Moving forward, the virtual histological staining pipeline may be explored using additional tissue types stained with various special and immunohistochemical stains to determine specific pathologic processes.

## MATERIALS AND METHODS

### Lung tissue samples

Seven lung tissue samples were used as part of this study. To evaluate the proposed virtual staining pipeline, four samples (S1-S4) from patients who had been lung transplanted because of PH, in the form of idiopathic pulmonary arterial hypertension (IPAH), were used. The samples had been collected from the pathology biobank at Skåne University Hospital and had been imaged for a previous study ^40^. To assess the applicability of our approach to additional histological stains, a heart and lung sample from a wild-type mouse fetus (embryonic day 18.5), previously imaged as a control sample for another study ^41^, was used. Additionally, to further assess the generalizability and repeatability of our registration framework, an additional human lung tissue sample from a patient diagnosed with heritable pulmonary arterial hypertension (HPAH), also previously used in the study by Westöö *et al* ^40^, and a second mouse heart and lung sample from Gawlik *et al* ^41^ (Suppl. Fig. S2) were used. The study was approved by the Colorado Multiple Institutional Review Board and by the regional ethical review board in Lund, Sweden (Dnr 2017/597 and Dnr 2019-01769). The use of mouse tissue was approved by the Malmö/Lund (Sweden) Ethical Committee for Animal Research (ethical permit numbers M17319 and M278024).

### Synchrotron-based phase-contrast microCT image acquisition

Prior to histological sectioning and staining of the human lung samples, two regions within each sample were scanned with synchrotron-based phase-contrast microCT (PC*µ*CT). The imaging was conducted at the X02DA TOMCAT beamline of the Swiss Light Source at the Paul Scherrer Institut (Villigen, Switzerland), where the same imaging protocol as in a previous study was used ^42^. Each CT-reconstructed volume had approximate dimensions of 4.2 × 4.2 × 2.7 mm^3^ with an isotropic 1.63 µm voxel size. Additionally, PC*µ*CT data of the mouse sample was obtained at the ANATOMIX beamline of the SOLEIL synchrotron (Saint-Aubin, France) with dimensions of 7.2 × 7.2 × 3.6 mm^3^ and an isotropic 0.88 µm voxel size.

### Histology slides preparation

Following PC*µ*CT, the samples were serially cut into 3 µm-thick sections, and every 12^th^ section was subjected to histological Elastic van Gieson (EvG) staining. The stained sections were scanned at 0.5 µm/pixel resolution using an Aperio ScanScope digital slide scanner (Leica Microsystems, Wetzlar, Germany). The number of EvG-stained sections for S1, S2, S3, and S4 was 8, 9, 8, and 12, respectively. The two additional samples, the mouse heart and lung sample as well as the HPAH human lung sample, were also serially sectioned, stained with hematoxylin and eosin (H&E), and scanned at the same image resolution as the EvG-stained samples.

### Image registration

The 2D-3D multi-modal image registration pipeline was conceived by refining a number of well-established image processing techniques and optimizing them both for computational efficiency and robustness. An overall scheme is shown in Fig. S1 and can be divided into four steps:

In the first (preprocessing) step, the serial histological sections from each sample were co-registered with one another to account for slight rotations under which the sections were mounted on the glass slides. First, the whole slide images (WSI) were down-sampled to match the effective pixel size of the microCT images. Following that, intensity-based rigid registration in combination with a regular step gradient descent that iteratively optimizes the mean square error (MSE) was used for all WSI-s.

The second step involved estimating the approximate axial position of the WSIs relative to the microCT volumes in order to crop the WSIs to roughly match the field of view (FOV) of the latter. Using rapid visual inspection, we identified the relevant region on the top slice of the histological stack and applied the same cropping parameters to the remaining registered histological sections. The selected region was intentionally largerby approximately 2 mm on each sidethan the microCT acquisition area, which easily helped reduce the extensive FOV of the WSIs and produce data more suitable for the subsequent step. Since this process was only required for the top slice, it was relatively swift (less than 2 minutes per block).

The third step aimed at finding the (virtual) plane within the microCT volumes that matches the cropped WSI, i. e. identifying the exact tilt angle. For doing so, we applied the algorithm by Chicherova *et al*. ^27^ to inverted grayscale and median-filtered (kernel size = 3 x 3 pixels) WSI-s and microCT slices. This algorithm is based on extracting Speeded Up Robust Feature (SURF) descriptors ^43^ which are then matched using the second-nearest-neighbor-criteria ^44^ to establish pairs between the two imaging modalities. Finally, RANSAC plane fitting ^45^ was used to fit a plane to the 3D point cloud of matching points and the resulting planes were virtually cut from the microCT volume.

The fourth and last registration step involved the precise 2D alignment of the cropped WSI-s with its corresponding microCT counterparts. For this purpose, the images were first globally registered using a similarity transform found by extracting and matching SURF feature descriptors through the *M*-estimator sample consensus algorithm ^46^. After this initial global registration, the microCT image and its corresponding cropped WSI were divided into overlapping blocks of 256 × 256 pixels (with an overlapping amount of 128 pixels), and (local) rigid registration parameters based on SURF feature descriptors were computed for each block and then combined to obtain a non-rigid displacement field for the entire histological patch. This displacement field was then applied to the histology images to obtain accurately matched microCT-histology image pairs.

### Quantification of registration accuracy

Registration accuracy was assessed evaluating the Target Registration Error (TRE), which was calculated based on a set of fiducial points manually annotated in the microCT histology image pairs before and after registration. The TRE is defined as ^47^:

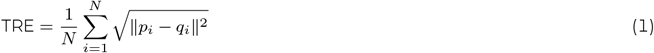

where *N* is the number of fiducial points, and *p*_*i*_ and *q*_*i*_ represent the coordinates of a fiducial point in the microCT and histology images, respectively.

For the TRE calculation, six microCT–histology image pairs were selected and eight fiducial points were manually annotated for each pair. To assess the generalization ability of the proposed registration pipeline, two of the image pairs were from one of the human lung samples with EvG-stained histological sections (S1), two from the human lung sample with H&E-stained histology images, and two from the mice heart sample with H&E-stained histologies. While a comparison of the proposed registration pipeline to other workflows is complicated since most are intended to either 2D or 3D registration (and not combined 2D-3D), to benchmark the proposed approach, we compared the quality of the alignment once the corresponding microCT plane to a given histology image had been found, using Elastix ^28^ - a widely used method for medical image registration -. Fiducial points were annotated on Elastix-registered images, and the TRE was also computed on those.

### Image preprocessing

Two types of artifacts needed to be accounted for before virtual histological staining, as discussed hereinafter. The histological images, prior to network training, were color-normalized using the *Color Transfer between Images* algorithm ^48^. This was to account for significant variations in color, illumination, and staining quality due to slight discrepancies in staining protocols and color responses of digital scanners ^49^. These may occur even when originating from the same tissue and stain. The microCT images, on the other hand, were background-corrected using a morphological image erosion linear plane-fitting algorithm ^50^. This was to correct the superimposed background intensity gradient, which is a typical artifact associated with local tomography ^51^.

### Virtual staining

For the virtual histological staining of microCT volumetric image data, we explored three different deep neural network architectures, of which the *pix2pix* image-to-image translation framework ^29^, based on a conditional Generative Adversarial Network (cGAN), was selected as the most suitable one. While indeed, to the best of our knowledge, this strategy represents the first successful application of *pix2pix* to histological and high-resolution microCT data, we also tested a(n) (unconditional) GAN, both as originally reported in a previous work on virtual histological staining of unstained tissue sections ^20^ as well as by reformulating the colorization as a pure regression problem by means of a fully convolutional network (FCN). All three approaches are briefly summarized as follows.

The final chosen model being based on a cGAN, comprises two networks: a U-Net-like generator that aims to learn the statistical transformation between the microCT images and their corresponding stained histologies, and a PatchGAN discriminator that learns to classify whether a given image is a real histology or, on the contrary, a virtually stained microCT image. In mathematical terms, the generator in a cGAN represents a mapping from a source domain image **y** ∼ *p*_source_ to its corresponding target domain image **x** ∼ *p*_target_ by estimating a mapping function 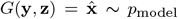, where **z** is a noise term. The cGAN objective function can be written as:

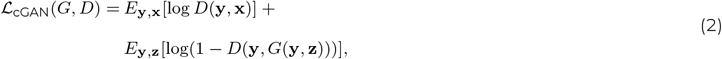

where the generator (*G*) minimizes the objective against the adversarial discriminator (*D*) that maximizes it. In addition, *pix2pix* includes the ℒ_*L*1_ pixel reconstruction loss to ensure a similar global structure between the output and ground truth image:

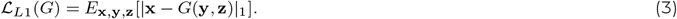

Putting all terms together, the final objective of the model can be written as:

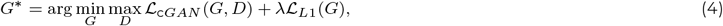

where *λ* balances both objectives terms and was set to 100 in our case.

The specific configurations of the generator and discriminator networks are illustrated in Fig. S3. In short, the generator consists of eight encoding and eight decoding blocks. Each encoder block comprises a 2D convolutional layer followed by a batch normalization layer and a leaky rectified linear unit activation. Each decoder block consists of a 2D transposed convolution, followed by a batch normalization layer, and a leaky rectified linear unit activation, except for the last layer in which a hyperbolic tangent activation function is used. In addition, skip connections are used to pass data between layers of the same level. The PatchGAN discriminator is conformed by four encoding blocks (the first one without batch normalization) followed by a 2D convolutional layer to map to a 1D output, followed by a sigmoid function. The PatchGAN made the real/fake classification after concatenating the input microCT image with the output of the generator or the ground truth histology images for ‘fake’ or ‘real’ examples, respectively.

Prior to the selection of this architecture, we evaluated the a(n) (unconditional) GAN model proposed by Y. Riverson *et al*. ^20^. This model again involves the training of two networks, a generator (*G*) and a discriminator (*D*). Both are optimized by minimizing the following losses:

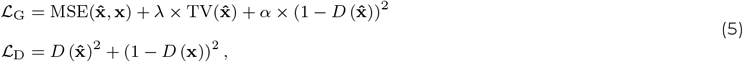

where **x** was the real histology and 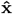 the virtually stained microCT images, respectively, with the hyperparameters *λ* and *α*. The main difference to before is that the discriminator makes the real/fake decision purely based on the histology output, thus omitting the concatenation of the microCT and histology images. The generator loss balances three terms: the mean square error (MSE), the total variation (TV) of the output image and the discriminator prediction of the output image. In their original work, the hyperparameters were chosen such that the TV loss accounts for 2% of the MSE, and the discriminator loss for 20% of the combined generator loss, respectively. In addition to this setting, we conducted a hyperparameter search and tested the following combinations for (*α, λ*): (19.000, 1), (30.000, 1), (47.000, 1), (1.000.000, 1), (30.000.000, 1). As we reported earlier, none of these combinations reached convergence during training. We therefore implemented six different image preprocessing steps aimed at bringing the microCT and histology images closer together in terms of appearance. The underlying hypothesis was that differences (such as noise distribution, ring artifacts, and resolved structures) between both imaging modalities prevented convergence from being achieved. The pre-processing settings are summarized in Table S1.

Finally, the above model was reformulated by omitting the discriminator and by introducing a weighted mean square error (wMSE) loss function as follows:

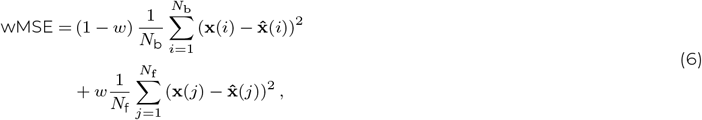

were *N*_b_ is the number of background pixels in an image, *N*_f_ is the number of foreground pixels, **x** denotes the physically stained histology image, 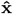 the virtually stained microCT image, and *w* is a scalar in the range [0, 1], representing the contribution of foreground pixels to the loss. This formalism resembled a fully convolutional network (FCN), which poses the colorization of microCT images as a pure regression problem, and has been widely used for style transfer applications in medical imaging prior to the emergence of GANs ^31,32^. Since, these types of networks have been extensively studied up to now, it served as an additional benchmark to the aforementioned two methods. In all our calculations, *w* was set to 0.7, to emphasize the accurate colorization of the lung tissue instead of the background.

### Implementation details

Before training, the microCT images were normalized to the intensity range [0, 1] and any intrusive air bubbles originating from the paraffin embedding process were masked out from them by setting those pixels to 0 intensity. For training the networks, the images (with dimensions 2560 × 2560 pixels in the EvG dataset and 4048 × 4048 in the H&E dataset) were cropped into patches of 512 x 512 pixels. To consider an image pair for either training or validation, we set a threshold for accepting only those image pairs for further processing when the correlation value between the microCT image patch and the corresponding histology was equal or greater than 0.45. This condition ensured that the network only learned genuine signals rather than perturbations caused by misalignments between both imaging modalities. Furthermore, only patches with 70% of the pixels contained within the inscribed circle of the microCT volume were considered, since only those pixels are correctly reconstructed after microCT phase-contrast retrieval. The image patches that satisfied all these conditions were then split into non-overlapping train (90%) and test (10%) sets, resulting in 536/60 images for the train/test sets of the EvG-virtual staining models, and 2610/290 for the H&E-virtual staining model. In the EvG cohort, image augmentation techniquesincluding rotation, flipping, and zoomingwere applied to patches containing blood vessels and respiratory airways to increase their representation relative to lung parenchyma and achieve a more balanced dataset.

Our virtual staining cGAN network was trained according to default parameters ^29^. The model was trained for 200 epochs with Adam optimizer and an initial learning rate of 0.0002, which was kept constant during the first 100 epochs and set to linearly decay to 0 during the last 100 epochs. The best generator (epoch 150) model was chosen manually by visually comparing the models every 10 epochs. The GAN model was trained using the optimizer, hyperparameters, and learning rates from the original implementation ^20^. The FCN was trained for 1, 000 epochs using the weighted adaptive moment estimation optimizer (AdamW) ^52^ with a weight decay of 10^−5^ and a fixed learning rate of 10^−4^. The last epoch was used for inference. The batch size for the networks was set to 10. The models were implemented using Python version 3.8.0, Pytorch version 1.9.0, and were trained on a computer with a Intel (R) Xeon (R) Gold 6248R CPU at 3 GHz and 64 GB of RAM, running a Red Hat Enterprise Linux 7.9 operating system, and using two Nvidia Tesla V100S GPU cards (each with 32GB GPU memory). Once the networks were trained, inference was performed on all axial slices of the microCT volumes.

### Evaluation metrics

Besides visual inspection and confirmation of correctly stained biological structures in the virtually stained microCT images by two board-certified pulmonary pathologists (CG, HB), we quantitatively evaluated the results using common metrics employed in virtual staining applications when paired output and ground truth images are available ^19^: Mean Squared Error (MSE), Structural Similarity Index Metric (SSIM) ^53^, and PSNR (Peak Signa-to-Noise Ratio) ^54^. Additionally, we considered LPIPS ^33^, as it has been reported to outperform previous metrics in terms of closeness to human perception of image quality. Following previous work ^55^, and to make the spatial frequency range identical between PCµCT and histology, we performed low-pass filtering (gaussian smoothing) with the same kernel size (*σ* = 2) for ground truth and VISTACT-generated images before metric calculations.

The MSE is defined as:

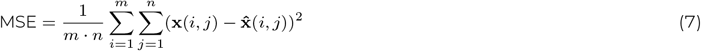

where **x** is the ground truth histology, 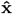 is the virtually stained microCT image (both normalized to the range [0,1]), and *m* and *n* are the number of pixels in the height and width image dimensions, respectively. The SSIM is based on the computation of three terms, namely the luminance term (*l*), the contrast term (*c*), and the structural term (*s*). The overall index is a multiplicative combination of the three terms, resulting in the following expression:

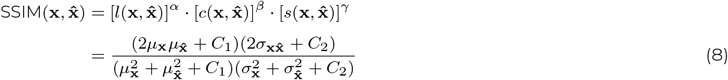

where 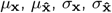, and 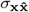 refer to the local means, standard deviations, and cross-covariance for images **x** and 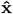. The default exponents are *α* = *β* = *γ* = 1, and the constants *C*_1_ and *C*_2_*/*2 were set to 6.5025 and 58.5225, respectively, by default. From the above equation, we observe that SSIM ranges between 0 and 1, with higher values indicating a greater similarity between the physically and virtually stained images.

PSNR is defined as:

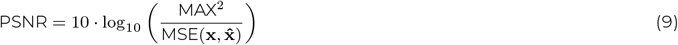

where MAX is the maximum possible pixel value of the image **x**. For example, for an 8-bit image, MAX would be 255. A higher PSNR value indicates better image quality.

Finally, the LPIPs metric encodes the ground truth and virtually stained histology using AlexNet ^56^ and computes the distance between their feature embeddings. For this metric, lower values mean higher similarity.

### Collagen segmentation

Collagen was automatically segmented from the virtually EvG-stained microCT volumes using color deconvolution ^37^. This technique involves an orthonormal transform of the color images, resulting in individual grayscale images corresponding to the concentrations of each employed stain. Subsequently, the resulting collagen channel was thresholded to generate a binary segmentation mask and image dilation with a 3-pixel-radius disk structural element was used for refinement purposes. All axial slices of a virtually stained microCT volume were then processed to obtain its corresponding 3D collagen segmentation mask. To ensure consistency in the segmentation across 2D axial slices, a 3-pixel kernel median filter was applied in the *Z* image dimension. This correction addressed potential false positives and false negatives not present in contiguous axial slices. Finally, the results from the segmentation pipeline were compared with masks obtained from the unstained microCT images using a semi-automatic machine learning model from Ilastik software ^34^. The segmentation was evaluated using the following pixel-wise metrics: Dice Similarity Coefficient (DSC), recall, precision, and accuracy, which were calculated based on the number of true positive (*T*_P_), false positive (*F*_P_), and false negative (*F*_N_) pixels, as follows:

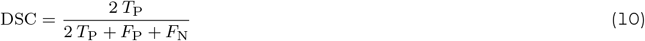

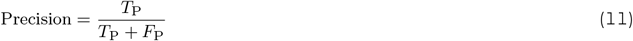

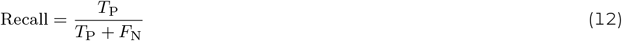

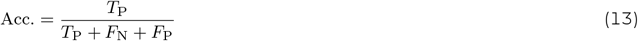

## Data and code availability

The PCµCT volumes and histology data along with the code (Github repository) used in this study will be made available upon publication acceptance.

## ACKNOWLEDGMENTS

C.A.-P. would like to acknowledge “La Caixa” Foundation (ID 100010434) under the fellowship code LCF/BQ/EU21/11890129 for enabling this project. We acknowledge the Paul Scherrer Institute, Villigen, Switzerland, for provision of synchrotron radiation beam-time at the X02DA TOMCAT beamline of the Swiss Light Source. We also acknowledge SOLEIL for provision of synchrotron radiation facilities, and we would like to thank Timm Weitkamp and Jonathan Perrin for assistance in using beamline ANATOMIX. We likewise acknowledge Martin Bech, Bodil Ohlsson, Robin Krüger and Julia Rogalinski for allowing and helping us to use some beamtime of SOLEIL proposal number 20220408 for this project. Schematic figures were created with BioRender.com.

## AUTHOR CONTRIBUTIONS

G.L., K.T.-L., and C.A.-P. designed research; C.A.-P. performed research and contributed analytic tools with input from G.L., N.P., K.G., K.T.-L. and A.H.S.; C.A.-P, G.L., N.P., K.T.-L., C.G. and H.B. analyzed data; M.S. contributed resources; C.A.-P. and G.L. wrote the paper with input from all authors.

