## Supplementary material for "Histology-guided 3D virtual staining of microCT-imaged lung tissue via deep learning"

### Figures

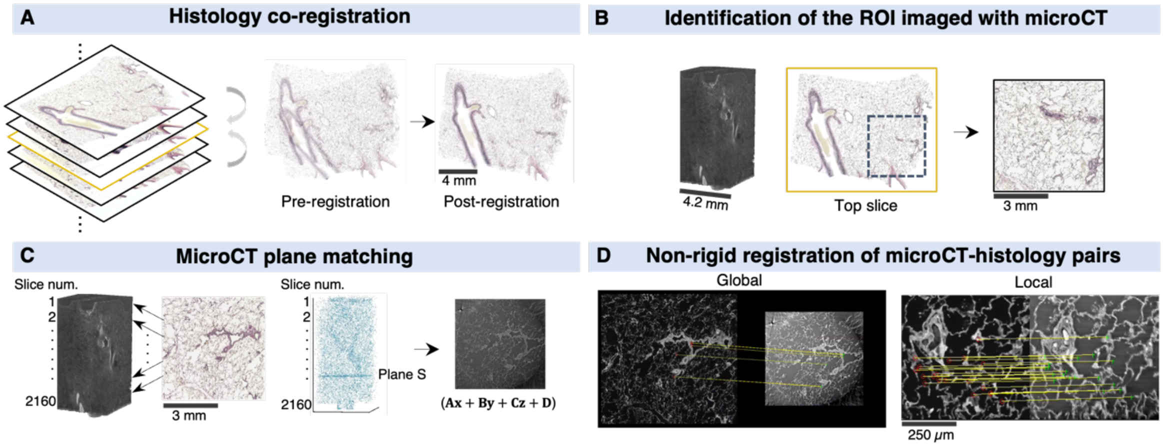


**Fig. S1. 3D – 2D multimodal image registration pipeline for microCT - histology image alignment. A.** Rigid intensity-based co-registration of serially cut histological sections. The central histology image was chosen as the reference for registration (fixed image). **B.** Manual identification of the approximate region of interest (ROI) from where microCT had been acquired. Identification was performed on the top section and applied to the whole (registered) histological stack. The identified region was ~2mm larger on each size compared to the acquired microCT. **C.** The oblique microCT plane corresponding to a histological ROI was found extracting SURF feature descriptors and matching those between the microCT volume and the histologic images. The matched points are shown in blue. RANSAC plane fitting in the matched points determined the corresponding microCT plane. **D.** Feature-based registration of the microCT-histology image pairs. Rigid global registration using SURF descriptors was followed by local registration in which the registration transformation was calculated on overlapping patches based on the local SURF descriptors. The registration was calculated on grayscale versions of the histologic images.

A

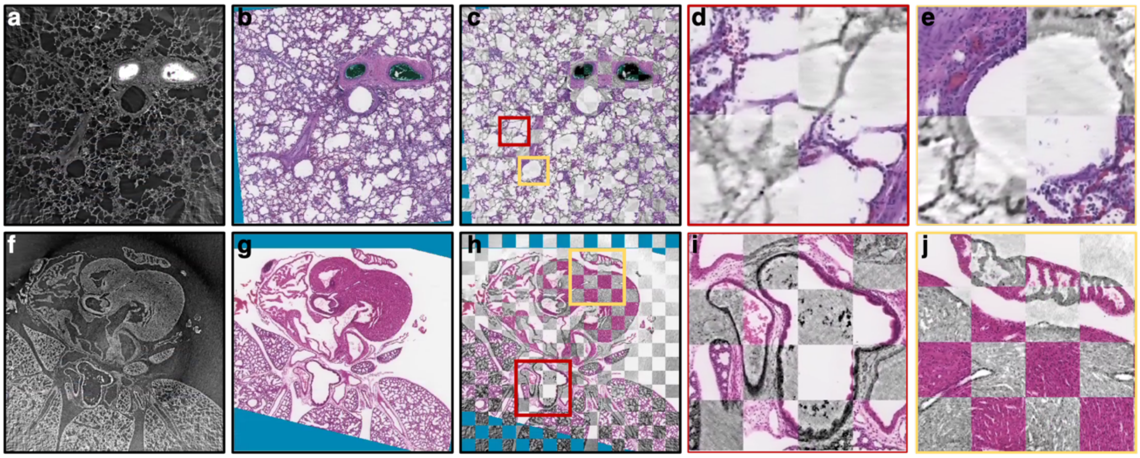


B

| | TRE (Prior reg.) | TRE (After reg.) |
|---|---|---|
| Lung dataset (H&E) | 0.13 ± 0.06 | 0.03 ± 0.04 |
| Lung dataset (EvG) | 0.60 ± 0.22 | 0.02 ± 0.02 |
| Heart dataset (H&E) | 0.52 ± 0.19 | 0.02 ± 0.03 |

C

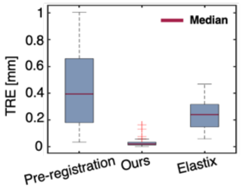


D

| Registration step | Time (min.) |
|---|---|
| Step 1: Histology co-registration | 1.71 |
| Step 2: ROI identification | 2.00 |
| Step 3: MicroCT plane matching | 55.30 |
| Step 4: Image pair alignment | 1.98 |

**Fig. S2. Qualitative and quantitative evaluation of the registration approach for employed for microCT-histology image alignment. A.** Visual inspection of the registration performance in a human lung sample (top row) and a mouse heart and lung sample (bottom row). **a, f** Phase-contrast microCT plane found with registration corresponding to the histological sections (b, g). c, h Overlay of the registered histology and microCT images using a checkerboard pattern. Intensity of the microCT image is shown inverted for clear visualization. **d, e, i, j** Zoom-in regions showing registered human alveoli (**d**), bronchiole (**e**), respiratory airway and accompanying vessel (**i**), and heart atrium and myocardium (**j**). **B.** Target registration error (TRE) once the corresponding 2D microCT plane to a given histology was found (prior reg.) and after the pairs of microCT-histology were aligned using SURF-based global and local registration (step 4 registration in *Methods,* after reg.). **C.** Boxplot of the TRE before the step 4 in the registration pipeline (pre-registration), after alignment with the proposed strategy (ours), and using Elastix for comparison purposes. **D.** Processing time estimates for each registration step. The values shown were calculated based on the alignment of the 8 EvG-stained histological sections of sample S1 to one of the microCT volumes of this sample.

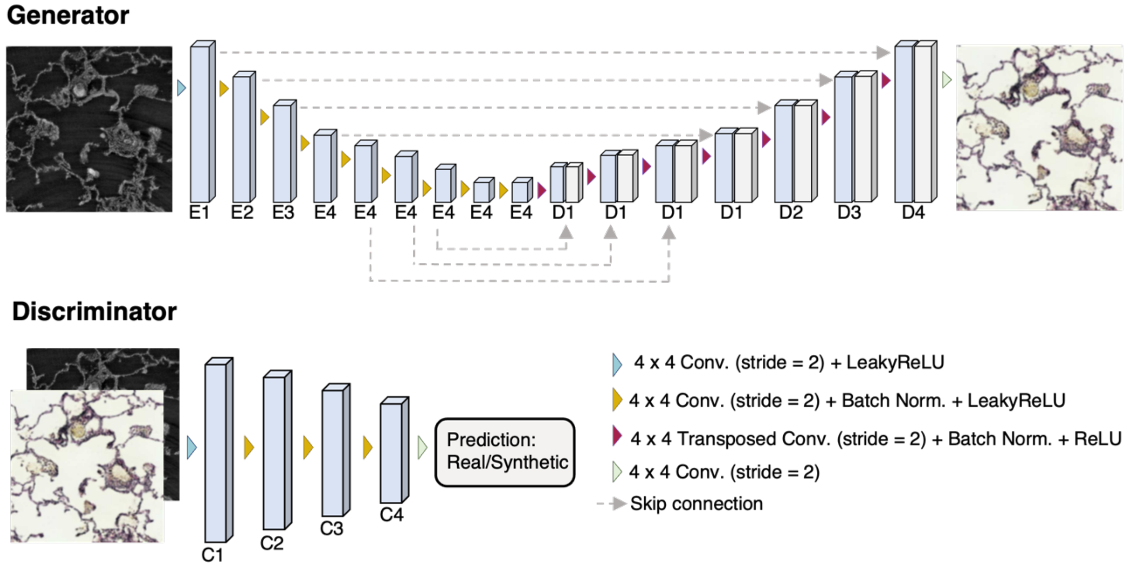


**Fig. S3. Conditional Generative Adversarial pix2pix model architecture.**The generator section consists of eight encoding and eight decoding blocks. Each encoder block comprises a 2D convolutional layer followed by a batch normalization layer and a leaky rectified linear unit activation. Each decoder block consists of a 2D transposed convolution, followed by a batch normalization layer, and a leaky rectified linear unit activation, except for the last layer in which a hyperbolic tangent activation function is used. Skip connections are used to pass data between layers of the same level. The number of channels at each stage of the encoder are E1=64, E2=128, E3=256, E4=512, and in the decoder D1=1024, D2=512, D3=256, and D4=128. The PatchGAN discriminator consists of four encoding blocks (the first one without batch normalization) followed by a 2D convolutional layer to map to a 1-dimensional output, followed by a sigmoid function. The number of channels at each stage is denoted by C1, C2, C3, and C4 and are 64, 128, 256, and 512, respectively.

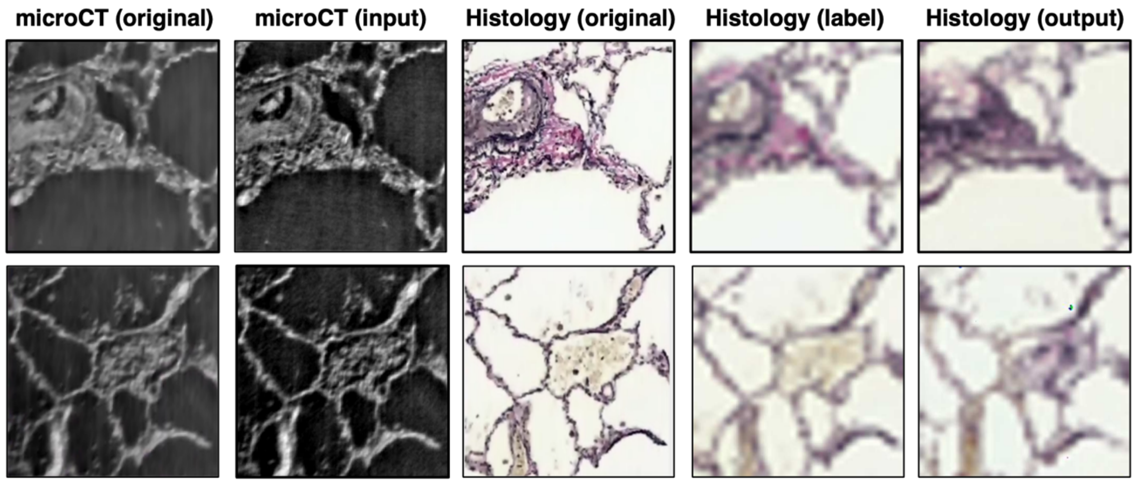


**Fig. S4. Virtual histological staining in pre-processed images with GAN model.** The preprocessing described in the last row of Table S1 was used. From left to right in each row: original microCT validation image, preprocessed microCT image with unsharp masking (input to the model), corresponding EvG-stained histology image, preprocessed histology image downscaled and upscaled by a factor of 8 (serving as the ground truth histology for the model), and lastly, the output image of the network.

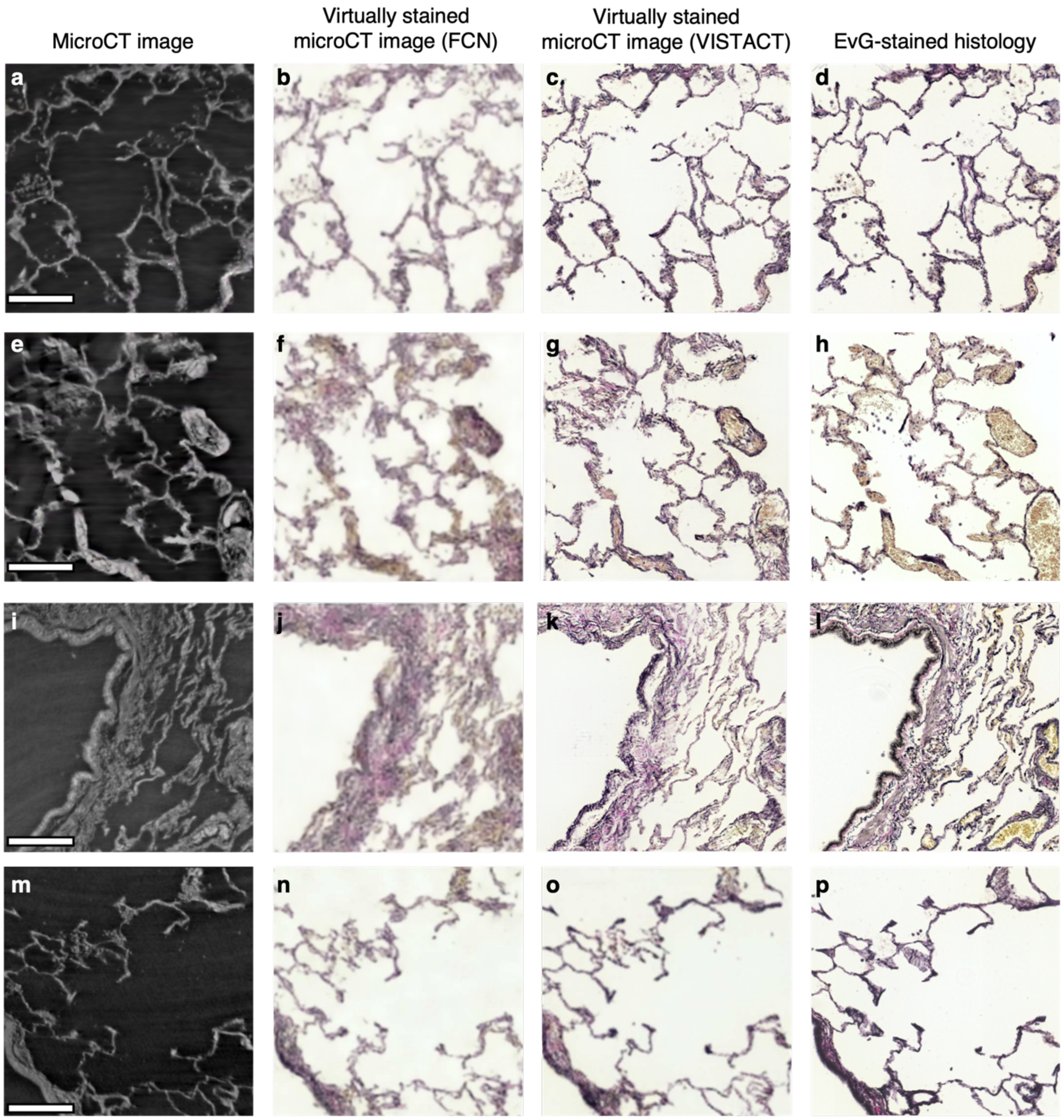


**Fig. S5. Qualitative comparison of FCN and cGAN (VISTACT) models for microCT virtual staining. a, e, i, m** Phase-contrast microCT images of human lung tissue used as input into the neural networks. **b, f, j, n** Virtually EvG-stained microCT images using a fully convolutional network (FCN). **c, g, k,o** Virtually EvG-stained microCT images with VISTACT. **d, h, i, p** Bright-field images of EvG-stained histological sections (ground truth for virtual staining). All scalebars are 200 µm.

**Tables**

**Table S1. Preprocessing techniques used to analyse the lack of convergence with GAN model.** Six image preprocessing pipelines aimed at bringing the microCT and histologic images closer together in terms of appearance were tested. The underlying hypothesis was that the differences (such as noise distribution, ring artefacts, and resolved structures) between both modalities were preventing convergence from being achieved. In unsharp masking, *σ* refers to the standard deviation of the Gaussian lowpass filter while *γ* refers to the strength of the sharpening effect. The parameter *r* of the median filter indicates the size of the neighborhood in the height and width image dimensions. Finally, in the last two rows, *f* denotes the resizing factor. Convergence was only achieved with the last preprocessing pipeline (bottom row).

| Histology preprocessing | MicroCT preprocessing |
|---|---|
| Gaussian filter (σ = 2 pixels) | Unsharp masking (σ, γ) = (10, 0.8) |
| Gaussian filter (σ = 2 pixels) | Unsharp masking (σ, γ) = (20, 0.8) |
| Median filter (*r* = 5 pixels) | Unsharp masking (σ, γ) = (20, 0.8) |
| Median filter (*r* = 7 pixels) | Unsharp masking (σ, γ) = (20, 0.8) |
| Downsampling followed by upsampling (*f* = 4) | Unsharp masking (σ, γ) = (20, 0.8) |
| Downsampling followed by upsampling (*f* = 8) | Unsharp masking (σ, γ) = (20, 0.8) |

**Table S2. Quantitative comparison of the FCN and cGAN (VISTACT) models for microCT virtual staining.** Average ± standard deviation of the evaluation metrics for images in the validation set.

| | MSE (↓) | SSIM (↑) | PSNR (↑) | LPIPS (↓) |
|---|---|---|---|---|
| **FCN** | **0.018 ± 0.006** | 0.62 ± 0.11 | **17.56 ± 1.46** | 0.30 ± 0.06 |
| **VISTACT** | 0.020 ± 0.007 | **0.67 ± 0.12** | 17.50 ± 1.63 | **0.25 ± 0.07** |

**Table S3. Segmentation metrics obtained with virtual histological staining (VISTACT) followed by image preprocessing and Ilastik.**

| | DSC (↑) | Precision (↑) | Recall (↑) | Accuracy (↑) |
|---|---|---|---|---|
| **VISTACT** | **0.65 ± 0.01** | **0.65 ± 0.05** | 0.66 ± 0.03 | **0.49 ± 0.02** |
| **Ilastik** | 0.47 ± 0.03 | 0.32 ± 0.03 | **0.87 ± 0.03** | 0.31 ± 0.03 |

**Table S4.** Quantitative performance of microCT H&E-virtual staining based on Mean Square Error (MSE), Structural Similarity Index Metric (SSIM), Peak Signal-to-Noise Ratio (PSNR) and Learned Perceptual Image Patch Similarity (LPIPs). Mean ± standard deviation of the evaluation metrics for images in the validation set.

| MSE (↓) | SSIM (↑) | PSNR (↑) | LPIPS (↓) |
|---|---|---|---|
| **0.0137 ± 0.0098** | **0.62 ± 0.22** | **21.61 ± 7.10** | **0.33 ± 0.14** |
